# Decision-making through integration of sensory evidence at prolonged timescales

**DOI:** 10.1101/385989

**Authors:** Michael L. Waskom, Roozbeh Kiani

## Abstract

When multiple pieces of information bear on a decision, the best approach is to combine the evidence provided by each one. Evidence integration models formalize the computations underlying this process [1–3], explain human perceptual discrimination behavior [4–11], and correspond to neuronal responses elicited by discrimination tasks [12–17]. These findings indicate that evidence integration is key to understanding the neural basis of decision-making [18–21]. Evidence integration has most often been studied with simple tasks that limit the timescale of deliberation to hundreds of milliseconds, but many natural decisions unfold over much longer durations. Because neural network models imply acute limitations on the timescale of evidence integration [22–26], it is unknown whether current computational insights can generalize beyond rapid judgments. Here, we introduce a new psychophysical task and report model-based analyses of human behavior that demonstrate evidence integration at long timescales. Our task requires probabilistic inference using brief samples of visual evidence that are separated in time by long and unpredictable gaps. We show through several quantitative assays how decision-making can approximate a normative integration process that extends over tens of seconds without accruing significant memory leak or noise. These results support the generalization of evidence integration models to a broader class of behaviors while posing new challenges for models of how these computations are implemented in biological networks.

## Results

The normative basis of the evidence integration framework and its ability to explain perceptual discrimination behavior suggest that its principles are broadly relevant to understanding decision-making. But at present, there are important obstacles to this generalization. While perceptual discrimination tasks afford tight experimental control and embody many important aspects of decision-making, they rarely demand integration at timescales that are characteristic of more complex natural behaviors. Humans can deliberate about natural decisions for many seconds (or even much longer) and, while doing so, often consider multiple discrete pieces of information that bear on their choice. The disparity between experimental tasks and natural behavior limits the application of current insights about perceptual discriminations to other domains. In particular, it might be expected that evidence integration would have limited broader relevance because the long and discontinuous timescale of evidence availability in the natural world would prove challenging for the neural mechanisms thought to underlie integration in rapid discrimination tasks.

To address this concern, we developed a new psychophysical paradigm that permits evaluation of evidence integration models in the context of prolonged decision-making (Figure 1). We built on the success of established perceptual discrimination tasks by providing evidence to the subject in the form of simple visual stimuli that could be parametrically controlled with high precision and whose basic encoding and representation in sensory cortex are well understood. But rather than displaying these stimuli in a continuous stream, we instead presented multiple brief exposures with variable intensity that were separated in time by long and unpredictable gaps. This simple manipulation produced a long and variable timescale of deliberation, extending it on many trials to tens of seconds (mean trial duration, 10.1±5.6 s; range, 2.2–34 s). We leveraged our experimental control over the sensory evidence and the variability in its strength, quantity, and timing to identify the computations underlying decision-making behavior.

We focused our analyses on three questions about the decision-making process. First, did it combine information from multiple samples before reaching a commitment? Second, did it use the graded quantity of evidence provided by each sample? Third, was it limited by memory leak or noise? Together, these questions allowed us to assess whether decisions could be made through evidence integration at prolonged timescales.

To answer these questions, we implemented a set of computational models that formalize different approaches for using the evidence to make a choice. These models generate quantitative predictions and exhibit distinct qualitative signatures that characterize different computational properties of the decision-making process (Figure S1). We focus here on four specific models: one implements the normative policy of evidence integration and serves as a baseline for comparison, while the others provide insight into each of the three questions enu-merated above. The models span a space of possible mechanisms within the general sequential samplingframe-work. They operate by using each sample, denoted x, to update a decision variable, denoted *V*. The evidence is quantified by the log-likelihood ratio (LLR) that the sample was generated from the high contrast distribution, and the sign of the decision variable at the end of the trial determines the choice.

**Figure 1:**
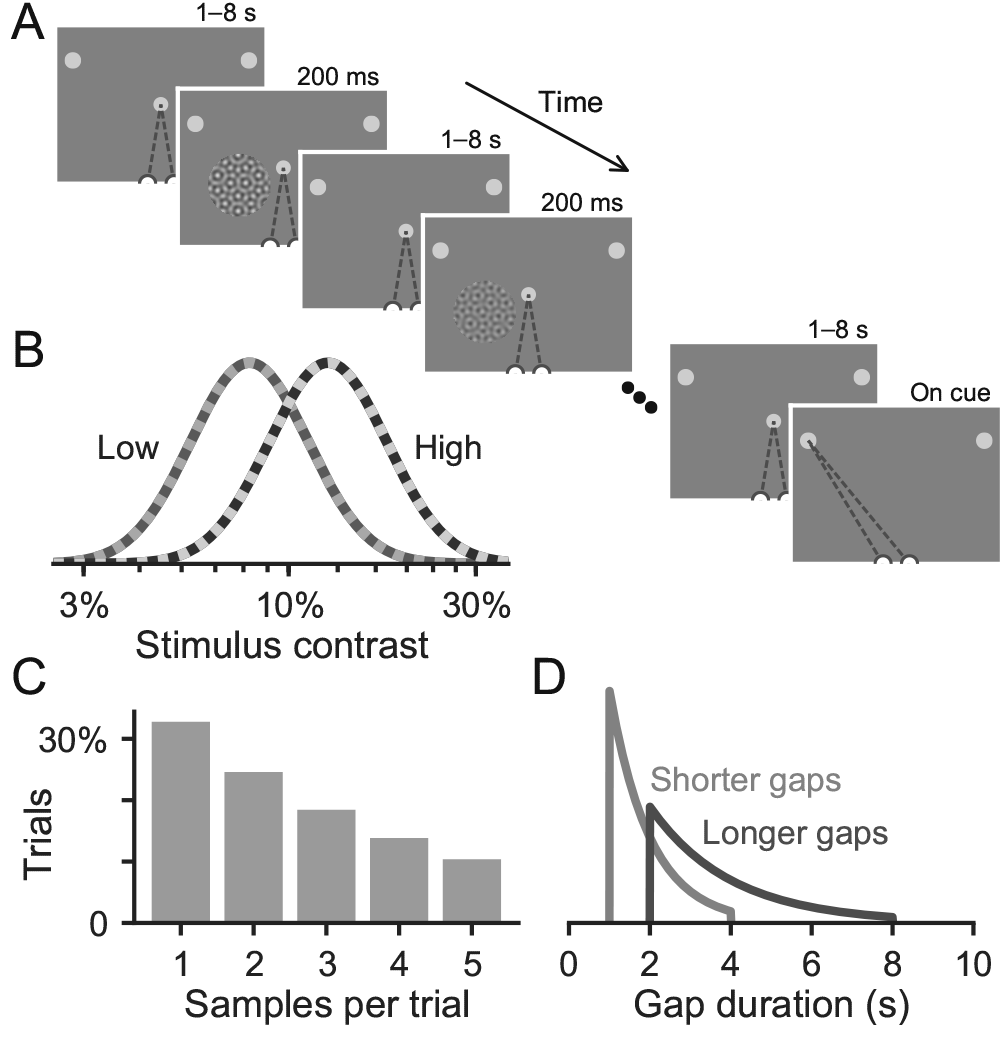
Experimental design. (A) Subjects viewed brief samples of a contrast pattern while maintaining central fixation. They were cued at the end of the trial to report their decision by making a saccade to one of two targets. (B) Each sample had a different contrast, randomly drawn from one of two overlapping Gaussian distributions in log contrast space. Each trial was generated using samples from the same distribution; the subject’s task was to infer which one. (C) 1–5 samples were shown before cuing a response, determined by drawing from a truncated geometric distribution. (D) Each sample was followed by a gap lasting either 1–4 s (shorter gap sessions) or 2–8 s (longer gap sessions), determined by drawing from one of two truncated exponential distributions.

### Integration of evidence across samples

Optimal performance in the task can be achieved by summing the evidence afforded by each sample in units of LLR with constant weighting across time. This computation can be formalized in a “Linear Integration” model defined by the following update equation:

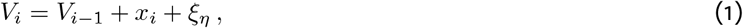

where *V_i_* is the decision variable after observing sample *i*, and *V*_0_ = 0. Choice variability arises in the Linear Integration model because the encoding of each sample is subject to Gaussian noise, represented by *ξ_η_* (Figure S1A). The only free parameter in the model is *σ_η_*, the standard deviation of *ξ_η_* in units of LLR. The decision variable updates are indexed by the ordinal sample number; this model does not use information about the time of sample appearance or the duration of the gaps. Linear Integration is optimal in that it is limited only by variability in the stimulus generation and noisy encoding of each sample; no other information is distorted or lost during deliberation [1–3].

Using this model, we derived analytic expressions for three assays of decision-making behavior (aggregate fits: Figure 2; individual subject fits: Figure S2; blue lines). Details of the mathematical derivations are provided in the methods section. The first behavioral assay shows how choice depends on the mean strength of evidence across samples (“sample mean psychometric function” or “mPMF”; Eq. 9; Figures 2A and S2A). It demonstrates that the sensory noise term in the Linear Integration model can account for behavioral variability. The second assay shows how accuracy depends on the number of samples in a trial (“sample count psychometric function” or “cPMF”; Eq. 10; Figures 2B and S2B). It demonstrates that performance benefits when more samples are shown, consistent with integration across samples. The third assay relates choice accuracy to stochastic fluctuations in stimulus evidence across time (“reverse correlation function” or “RCF”; Eq. 11; Figures 2C and S2C). It demonstrates approximately equal weighting of each sample. Each effect was replicated in individual subjects (Figure S2), suggesting that integration of evidence can explain both the average behavior and the idiosyncrasies of individual subjects.

An important point related to interpreting the RCF should be noted. Both the data and Linear Integration model exhibit an apparent nonlinearity resembling a primacy bias in the onset-aligned RCF and a recency bias in the choice-aligned RCF. But this nonlinearity is an artifact of averaging across trials with different sample counts, because the influence of each sample on choice scales inversely with the square root of the total number of samples [27]. The equal weighting of each sample can be seen by the close match between the data and the Linear Integration model, which has aflat RCF for each sample count (equation 16).

**Figure 2:**
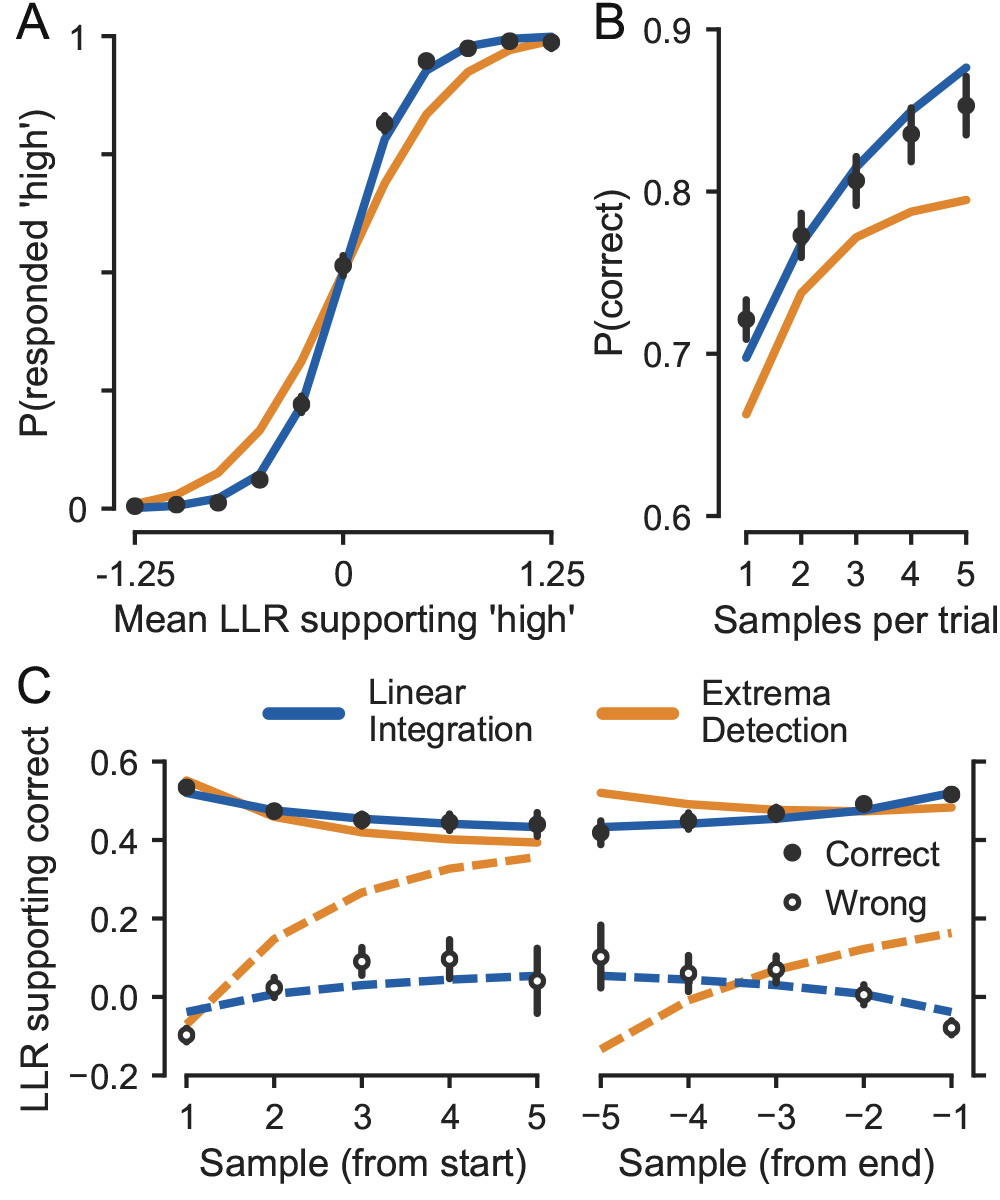
Integration of evidence across samples. (A) Sample mean psychometric function (mPMF), showing the relationship between mean evidence strength supporting a choice of “high” and choice probability. (B) Sample count psychometric function (cPMF), showing the relationship between the number of samples in a trial and the probability of making a correct choice. (C) Reverse correlation functions (RCFs), aligned to the start of the trial (left panel) or the end of the trial (right panel), showing the conditional mean of the evidence over time given choice accuracy. In all panels, black points and lines show means and bootstrap 95% CIs, and blue and gold lines show analytic functions from fitted Linear Integration and Extrema Detection models, respectively. All panels show aggregate data and model fits; see Figure S2 for individual data and fits.

Based on these three assays, the choice data appear generally consistent with evidence integration. Yet minor quantitative deviations from the Linear Integration model (e.g., a shallower cPMF) imply that it is not a full account of the decision-making process. While several different factors could account for minor deviations from normative performance (see Discussion), a primary concern is whether the data can be explained by a process that lacks integration altogether. The strongest candidate would be a model that performs decision-making through sequential sampling but without integration of evidence. This model compares each sample individually against a detection threshold; if the threshold is exceeded, the process terminates in a commitment to the corresponding alternative, and further samples are ignored. Prior to commitment, intermediate samples of evidence are discarded and have no bearing on choice. If no sample produces a commitment, then a response is generated randomly at the end of the trial. The process can be termed “Extrema Detection” [28] and leads to the following update equation:

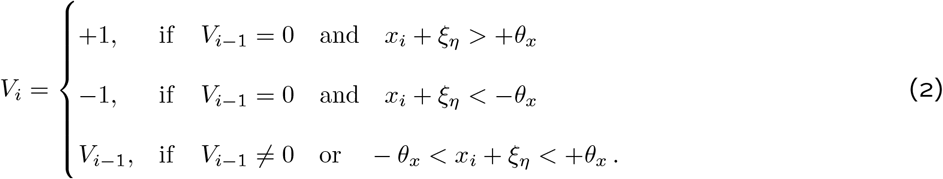

As in the Linear Integration model, *ξ_η_* represents Gaussian noise with standard deviation *σ_η_*; the other free parameter, *θ_x_*, represents the detection threshold.

Despite using fundamentally different computations, Extrema Detection can superficially mimic evidence integration: both the strength and number of samples will influence choice (Figure S1B). But fitting the model to the choice data shows that it cannot explain the subjects’ behavior. The Linear Integration model had a higher cross validated log-likelihood (aggregate Δ*LL_CV_*: 788.7; individual Δ*LL_CV_*: 95.2–288.4). The three behavioral assays also revealed a strong divergence between data and model (aggregate fits: Figure 2; individual fits: Figure S2; gold lines), further demonstrating the broad failure of the Extrema Detection model to account for qualitative features of decision-making. Therefore, the observed behavioral performance could not have been obtained using only individual samples, implying that evidence was integrated across the gaps.

### Integration of graded stimulus evidence

The normative model in Eq. 1 uses the graded weight of evidence afforded by each sample in units of LLR. Another source of deviation from normative integration could involve a transformation applied to the evidence values before they are combined across trials. One extreme form of transformation would be to binarize the evidence from each sample as supporting either a “high” or “low” choice. This operation would cause the decision variable to reflect a count of the number of samples supporting each choice rather than the integrated continuous evidence. The resulting model (“Counting”) can be defined with the following update equation, using the same terms as in Equation 1:

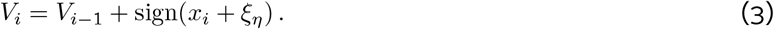

The Counting model is structurally similar to the Linear Integration model and makes broadly similar predictions (Figure S1C). When the model was fit to the choice data, the likelihood was only moderately decreased relative to Linear Integration (aggregate Δ*LL_CV_*: 71.5; individual Δ*LL_CV_*: −0.5–27.6). Despite this similarity, the Counting model exhibits a distinctive qualitative signature that is not present in the data. Representing the decision variable with a discrete count leads to a tie on many trials with an even number of samples, requiring a random guess to generate a choice. As a result, the model’s expected accuracy does not improve relative to trials with the next smallest odd number of samples, a prediction that is independent of the sensory noise parameter (Figure S1C; Equation 37). In contrast, the data exhibit a clear improvement in accuracy with additional samples (Figures 3A and S3B) and have statistically significant improvements on trials with even numbers of samples relative to those with the next smallest odd number (Eq. 6; aggregate model: *β*_1_ = 0.25±0.04; *P* < 1e-8; individual models: 5/5 subjects *P* <0.005).

**Figure 3:**
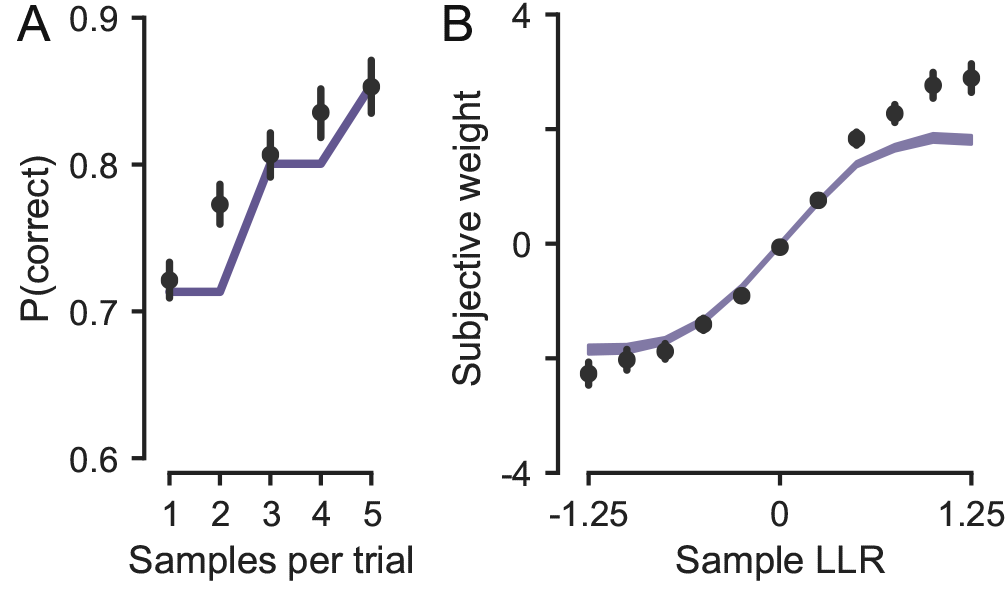
Integration of graded stimulus evidence. (A) Data and model cPMFs, showing how the data deviate from a qualitative signature of discrete evidence transformation. Black points and error bars show means and bootstrap 95% CIs; purple line shows analytic prediction of the Counting model. (B) Estimated subjective weighting of samples with different evidence values. Black points and lines show logistic regression coefficients and 95% CIs for the data; purple band shows logistic regression coefficients and 95% CIs for simulated data from the Counting model with parameters that best fit the choices. Both panels show aggregate data and model fits; see Figure S3 for individual data and fits.

The loss of information from a discrete transformation can also be evaluated by estimating the subjective weight of evidence afforded by samples with different objective strengths (Eq. 5). In the Counting model, discrete transformation curtails the influence of strong samples. By comparing the subjective weights estimated from the choice data to weights obtained by simulating choices from the best-fitting model (Figures 3B and S3C), it is apparent that strong samples contributed more to the decision than would be expected after a discrete transformation. Therefore, the data are most consistent with integration of graded stimulus evidence across multiple samples.

### Minimal influence of memory leak or noise

The normative model in Eq. 1 does not require any information about the timing of the sample presentations or the duration of the gaps between them. We expected, however, that long deliberation timescales would cause deviations from normative integration due to limitations in working memory. We reasoned that this might happen in two ways. First, information from earlier samples could fail to influence the subject’s choice if it “leaked” from memory [22]. Second, additional behavioral variability could emerge from integration of diffusive noise in the absence of sensory input [23]. To test for the presence of these limitations, we extended Eq. 1 into the temporal domain and explicitly modeled the influence of these two factors on the decision variable (Figure S1D). This “Leaky Integration” model can be defined by the following update equation:

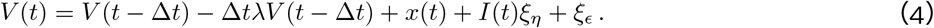

In this equation, λ represents the memory leak rate (the inverse of the integration time constant *τ*) in units of seconds. *x*(*t*) represents the strength of evidence at time *t*, if a sample was presented, and is 0 during the gaps. *I*(*t*) is an indicator variable that specifies when a sample is visible and governs the influence of sensory noise, 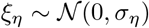. Finally, 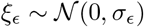 represents memory noise, which accumulates throughout the gaps.

Surprisingly, fitting the Leaky Integration model to choice behavior revealed minimal influence of these factors (Figures 4 and S4). After optimizing the free parameters to maximize the likelihood of the choices, we simulated datasets from the Leaky Integration model and compared them to the behavioral data and to the fits of the normative Linear Integration model. This comparison showed that Leaky Integration could not be distinguished from the normative Linear Integration model in terms of its predictions about the cPMF (Figures 4A and S4A) or RCF (Figures 4B and S4B). Correspondingly, the likelihood only marginally exceeded that for the Linear Integration model, despite the two additional free parameters (aggregate Δ*LL*_CV_: −5.1; individual Δ*LL*_CV_: −8×10^8^ – −5.1; aggregate likelihood ratio test: 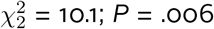; individual likelihood ratio tests *P* <0.05 in only 1 of 5 subjects). Both the optimized memory leak rates and memory noise magnitudes were close to zero (Figure 4C), implying an integration time constant longer than 25 s for all subjects and an effectively infinite integration time constant for 4/5 subjects.

The minimal influence of the long integration timescale can also be seen using a model-free approach. Subjects performed the task in two different conditions where the gaps were sampled from distributions with different minimum, maximum, and mean durations (Figure 1D). Despite these differences, performance on the task did not appear to vary substantially between the two conditions (Figures 4D and S4C). Statistically, behavioral sensitivity to the strength of evidence was similar between conditions (Eq. 7; aggregate model: *β*_3_ = 0.29±0.18; *P* = 0. 11; individual models: *P* > 0.05 in 4/5 subjects and *P* = 0.015 in S4). In fact, one could use the behavior in one of the conditions to predict the behavior in the other. These results further emphasize the functional invariance of the decision-making process to large differences in the timescale of evidence availability across trials, implying the existence of a highly flexible and efficient system for combining the evidence from each sample to make decisions at a level approaching normative performance.

**Figure 4:**
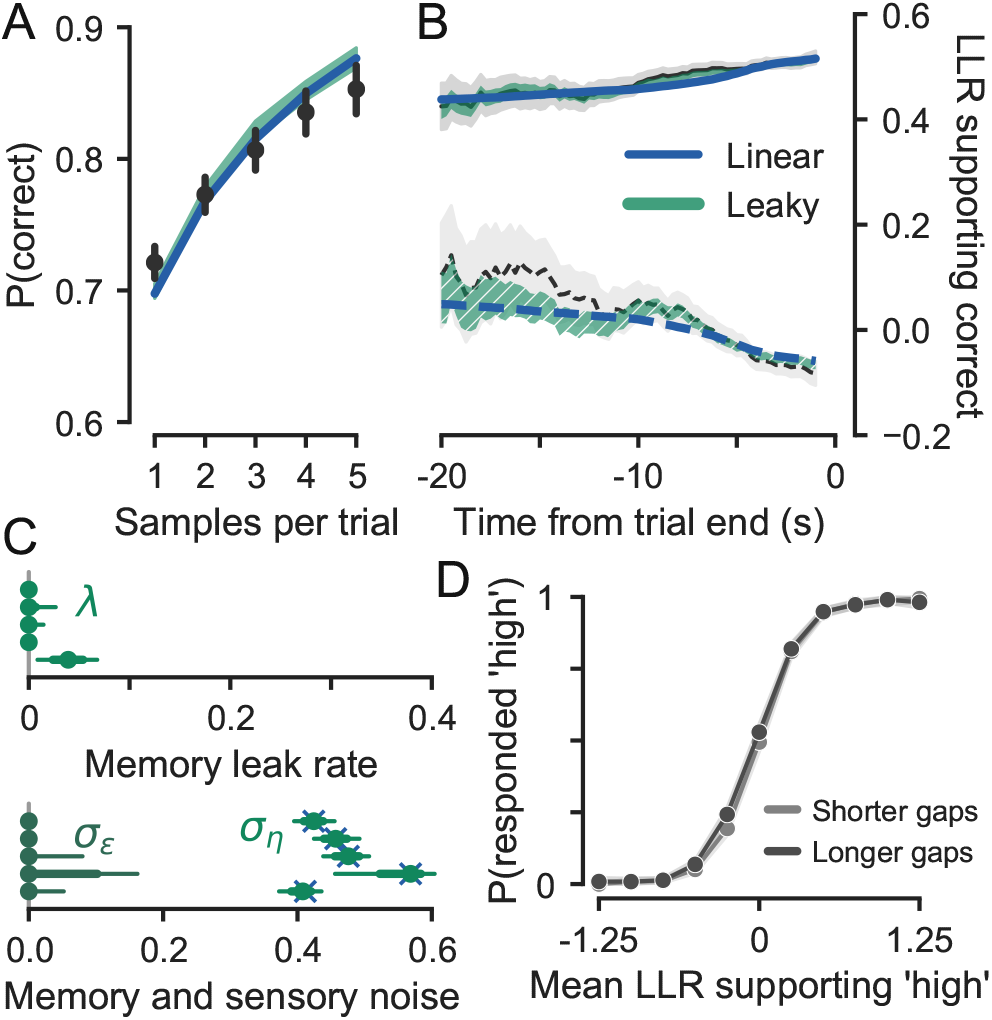
Minimal influence of memory leak or noise. (A) Data and model cPMFs. Black points and error bars show means and bootstrap 95% CIs for the data, blue line shows analytic function for the best-fitting Linear Integration model, and green band shows simulated performance for the best-fitting Leaky Integration model. (B) Data and model RCFs, aligned to the end of the trial. Element colors are as in panel A; solid lines show functions conditioned on correct choices and dashed lines show functions conditioned on errors. (C) Integration model parameter fits. Green elements show parameters for the Leaky Integration model: points show maximum likelihood estimates; thick and thin error bars show bootstrap 68% and 95% CIs, respectively. Blue crosses show *σ_η_* estimated with the Linear Integration model. (D) Data mPMF plotted separately for trials with shorter and longer gaps between samples. All panels show aggregate data and model fits; see Figure S4 for individual data and fits.

## Discussion

The role of evidence integration in perceptual decision-making has been most commonly studied in tasks where a subject engages with a sensory stimulus for a continuous and relatively brief amount of time. The introduction of these tasks represented an innovation permitting the study of mechanisms underlying deliberative processes on the timescale of hundreds of milliseconds [29]. By transcending simple reflexes while remaining accessible to rigorous computational analysis and invasive neural recordings, they contributed to development of a theory that spans multiple levels of analysis to explain how sensory information guides goal-directed choice [10, 30]. More recent innovations have extended this theory in important ways. The creation of pulsatile, count-based discrimination tasks provided additional leverage for detailed computational modeling [5, 15, 31] and facilitated neurophysiological investigations in new model systems [32–34]. Extending the framework to probabilistic inferences on the basis of rapid counting [35], arbitrary learned associations [16, 36], or parametric stimulus properties [37, 38] has contributed to further understanding of the decision-making process, including what quantities it operates on and why it might fail to achieve optimality. Yet there has been persistent concern about the scope of behaviors to which these insights apply.

Long timescales might be expected to pose a challenge for evidence integration because of biophysical limitations in neurons. Limitations of individual units can be surmounted through interactions in neural networks, but existing network models have been tested primarily in decision-making tasks demanding integration over much shorter timescales. These models propose that a careful balance between recurrent excitation and inhibition can facilitate the emergence of attractor dynamics, extending their intrinsic timescales beyond those of their constituent units [3, 22–24, 26, 39]. Such networks can implement evidence integration, but they will have failure modes that become apparent when pushed beyond their intrinsic timescales. Rapid discrimination tasks are often modeled by networks with bistable point attractor dynamics [40, 41]. In our task, these networks might be expected to implement Extrema Detection because, in the sustained absence of input, a decision variable computed with point attractor dynamics would either decay back to its starting point or converge onto a terminal state based on one sample. An alternate modeling framework uses line attractor dynamics to represent and manipulate graded quantities of evidence [42, 43]. But that representation would likely be corrupted by accumulated noise during prolonged gaps [23, 44], and reducing the influence of noise with memory leak would erase evidence from early samples [22].

That none of these failure modes emerged when we pushed the decision-making process an order of magnitude beyond the timescale of conventional tasks suggests either that the intrinsic timescale of biological networks can be much longer than currently thought [25, 26] or that implementation-level models of evidence integration require augmentation. One possible accommodation would involve additional mechanisms that add robustness to noise [45, 46]. Alternatively, prolonged integration may be achieved by a functionally feedforward network architecture [47]. It may also be the case that prolonged integration recruits fundamentally distinct neural mechanisms to perform functionally similar computations. Indeed, invariance to gap duration is perhaps more characteristic of a long-term memory system than of the working memory mechanisms that attractor networks model. Can evidence integration at long timescales be implemented through interactions between multiple memory systems? As this question comes into focus, we note that the prolonged yet controlled timescale of evidence in our task would make the integration process amenable to study with functional MRI, complementing previous efforts that have leveraged gradual yet continuous perceptual judgments [48, 49].

While behavior approximated the normative policy for our task, minor but systematic deviations from the model fits imply that Linear Integration cannot fully explain the data. The behavioral assays suggest two sources of deviation. First, Figure 3 indicates that the influence of strong individual samples was moderately curtailed. Down-weighting of outlying evidence values would be consistent with previous observations in rapid perceptual averaging tasks [50, 51]. It may also reflect complexities in contrast perception that our models do not account for [52]. Second, Figure 2 suggests relative over-weighting of early samples, consistent with the presence of a decision bound that terminates integration prior to the response cue. A termination bound is a fundamental component of the theory explaining rapid discriminations [7, 10, 14, 30], so this explanation would not undermine our main conclusions. We do not quantitatively evaluate this model because, despite the presence of these qualitative signatures, our specific experimental design parameters make the height of the decision bound difficult to estimate. Nevertheless, the asymmetry between the RCF for correct and error trials suggests that integration processes were not uniformly terminated as in a simple bounded integration model. It is possible that a decision bound enacts only provisional commitments that can be revised by further evidence [53], perhaps with some cost for doing so [54]. Our task involves many trials where an optimally-computed decision variable would need to change sign, so deviations from normative behavior might also be explained by a resistance to changes of mind, similar to confirmation biases that over-weight initial impressions [37, 55]. Variants of our task that accentuate the influence of these factors may afford computational modeling of their origin and provide insight into suboptimal decision-making in naturalistic contexts.

Our results demonstrate that neither memory leak nor memory noise pose a fundamental challenge for using evidence integration to make naturalistic decisions. While surprising in light of biophysical properties of neurons, these findings create an opportunity to generalize the powerful evidence integration framework beyond simple perceptual judgments. We used visual stimuli to retain precise control over the quantity of evidence available to the subject at each moment in time. But in principle, these computations could be applied to use any relevant information that can be converted to the common currency of a decision variable. Many open questions remain about the computational and neural basis of natural decision-making. Answering them will require formal models that can make connections across levels of analysis to explain how biological systems generate complex behaviors. Evidence integration is a promising candidate for these efforts.

## Methods

### Subjects and experimental apparatus

Five human subjects, (four male and one female; ages 19–40) participated in the experiment. All had normal or corrected-to-normal vision. One of the subjects was author RK; the others were naive to the purposes of the experiment. All experimental procedures were approved by the Institutional Review Board at New York University, and the subjects provided informed written consent before participating.

The subjects were seated in an adjustable chair in a semi-dark room with chin and forehead supported before a cathode ray tube display monitor (21 in Sony GDM-5402; 1600 × 1200 screen resolution; 75 Hz refresh rate; 8 bit color; 52 cm viewing distance). The display was calibrated with a photometer and gamma corrected to have a linear response. The mean luminance of the display during the experiment was 35 cd/m^2^. Viewing was binocular. Gaze position was monitored at 1 kHz using a high-speed infrared camera (SR-Research; Ottowa, Canada). Stimulus presentation was controlled using *PsychoPy* [56].

### Experimental design

The experimental task is diagrammed in Figure 1. Each trial was initiated when the subject looked at a central fixation point (0.3° diameter). 500 ms after fixation, two response targets (0.5° diameter) appeared 5° above and 10° to the left or right of fixation, and a small protuberance (0.1° length) extended from the fixation point to cue the location at which the visual stimulus would be presented on that trial.

The visual stimulus was a circular (6° diameter) contrast patch constructed as the average of 8 sine wave gratings at evenly spaced orientations. Each grating had a spacial frequency of 2 cycles per degree, and the phases of the gratings were independently randomized on each presentation. The stimulus edges (20% of the radius) were blended with the background using a raised-cosine mask. The stimulus was centered 2°below and 5.6° either to the left or right of fixation and was always compatible with the cue. Within a given trial, the stimulus would appear in the same location, but the location varied randomly between trials. Each stimulus presentation had a duration of 200 ms, and there were 1–5 presentations per trial (drawn from a truncated geometric distribution with *p* = 0.25).

The contrast of the stimulus varied across each presentation. Stimulus contrast, *C*, was defined in units of Michelson contrast and was determined by setting each constituent grating to a contrast of 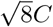 before averaging. This procedure generated contrast patches with root-mean-square (RMS) contrast that was equivalent to single gratings drawn at the specified value of *C*.

The experimental task required probabilistic inference on the basis of stimulus contrast. Each trial was defined, with equal probability, as either a “low-contrast” or “high-contrast” trial. This determined which of two overlapping distributions would be used to generate the contrast for each stimulus presentation in that trial. The distributions were defined as Gaussians in log_10_ contrast space with means of −1.1 or −0.9 and standard deviations of 0.15. The subject’s task was to infer, using the perceived contrast of each stimulus presentation, whether the trial was generated using the low-contrast or high-contrast distribution.

Each stimulus presentation was separated by a gap with a duration drawn from a truncated exponential distribution. The data reported here were collected across two different experimental conditions: “shorter” gaps (1–4 s) and “longer” gaps (2–8 s). These conditions appeared in separate experimental sessions, and subjects performed the shorter gap sessions first. The gaps prior to the first and following the last stimulus presentations were determined using the same distribution as the inter-stimulus gaps.

At the end of the trial, the fixation point disappeared, which cued the subject to report their decision by making a saccadic response to one of the two response targets. After choosing one of the targets, the subject received auditory and visual feedback about the accuracy of their response. Accuracy was defined in terms of whether the subject chose the target corresponding to the distribution that was actually used to generate the stimuli for that trial. Response target assignments were stable across the experiment. The subject was allowed to blink during the trial but was otherwise required to maintain fixation prior to responding. Trials were separated by an inter-trial interval of at least 3 seconds.

Each subject was trained on the task for 2–4 sessions prior to collection of the data reported here, and they were required to reach a criterion level of performance (session-wise accuracy >76%) before continuing. They additionally performed two practice sessions with longer gaps prior to collection of longer-gap data for analysis. Experimental instructions emphasized accuracy, and subjects were explicitly informed that the best way to maximize performance would be to make their decision on the basis of what they perceived as the average contrast of all the stimuli presented in each trial. Subjects typically performed 6 blocks (7 minute duration) in each session and received feedback about their accuracy after each block.

Not including training, each subject contributed 18 separate 1-hour sessions to the data reported here (6 sessions with shorter gaps and 12 sessions with longer gaps). In total, our analyses involved 14,869 trials (6,500 with shorter gaps and 8,369 with longer gaps).

### Data analysis

For our primary analysis of the different computational models, we characterized the data using three behavioral assays: one relating variability in behavioral responses to average stimulus strength (the sample mean psychometric function; mPMF), one relating choice accuracy to the number of evidence samples (the sample count psychometric function; cPMF), and one relating choice accuracy to stochastic variability in the stimulus over time (the reverse correlation function; RCF).

The mPMF illustrates how behavioral responses depended on the average strength of evidence in each trial. The evidence value corresponding to each stimulus contrast was defined using the log-likelihood ratio of the two generating distributions (positive evidence supports “high” and negative supports “low.”) Our experimental design produces a large amount of trial-to-trial variability in the strength of available evidence. To characterize the effect of this variability, we computed the proportion of “high” choices within evenly spaced bins of trials defined by the mean evidence across samples (bin width: 0.25, except for the lowest and highest bins which were unbounded on one side).

The cPMF illustrates how accuracy depended on the number of stimulus samples that were presented on each trial. To characterize this relationship, we computed the proportion of trials with the same number of samples where the subject chose the correct target (the target corresponding to the distribution that generated the observed evidence).

The RCF provides insight into the dynamics of the decision-making process by estimating the relative leverage on choice of evidence presented at different points in time either looking forward from the start of the trial or backwards from its end. This approach exploits the random variability in the strength of evidence afforded by each stimulus presentation. To estimate this function, we grouped trials based on response accuracy and then computed the mean strength of evidence supporting the correct target across time. Note that because we are conditioning on accuracy and not choice, the resulting function should not be considered an estimate of the sensory kernel. Instead, we are using the logic of reverse correlation to define a function that can be compared between data and model. This function represents a conditional expectation as a function of time. For some analyses, we considered the time dimension as ordinal (Figures 2C and S2C) and averaged over the first stimulus presentation of every trial, the second stimulus presentation of every trial with more than one sample, and so forth. For other analyses (Figures 4B and S4B), we used the actual timing of stimulus events by computing a running average over all stimuli occurring within a bin of width 4 s at evenly spaced timepoints (bin width: 0.25 s).

To estimate the influence of individual samples with different objective weights of evidence on the subjects’ choices, we used an approach described in previous work [36]. This analysis estimates the subjective weight of evidence afforded by samples falling into evenly spaced bins (bin spacing: 0.25) using the following logistic regression model:

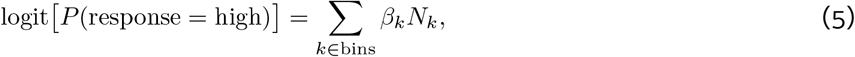

Where *N_k_* is the number of samples appearing in each trial with objective strength falling in bin *k*. We binned stimulus strengths because the objective sample strengths were continuous. The regression coefficient corresponding to each bin, *β_k_*, can be interpreted as an estimate of the subjective weight of samples falling within that bin.

Figures that show these functions for behavioral data have 95% bootstrap confidence intervals [57]. Confidence intervals were computed by resampling trials with replacement and estimating the statistic of interest across 10,000 iterations. The error bars correspond to the 2.5 and 97.5 percentiles of the resulting distribution.

We additionally performed several statistical analyses of choice behavior using logistic regression. To determine whether choice accuracy improved consistently with additional samples, we tested whether accuracy differed for trials with odd and even sample counts using the following function:

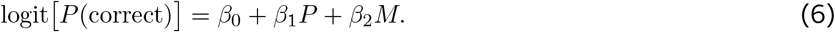

In this equation, *P* is an indicator variable specifying the parity of the sample count on each trial (odd vs. even), and *M* is an indicator variable specifying whether the sample count on each trial was 1–2 or 3–4. This analysis excluded trials with 5 samples because corresponding trials with 6 samples did not occur in the experiment. The null hypothesis (a prediction of the Counting model) is that sample parity would have no effect (*H*_0_ : *β*_1_ = 0).

To test for differences in behavioral performance between the longer and shorter gap conditions, we used a logistic regression model to test for an interaction between the influence of the mean evidence strength and the gap condition:

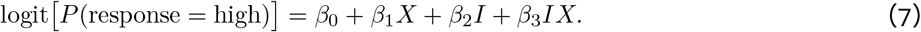

In this equation, *X* is the mean strength of evidence across samples in a trial, and *I* is an indicator variable specifying the gap duration condition. The null hypothesis was that the influence of stimulus evidence on choice did not vary across gap duration conditions (*H*_0_ : *β*_3_ = 0).

### General computational modeling approach

We identified the computational properties of the decision-making process by fitting and evaluating several quantitative models. The models spanned a space of computations within the general sequential sampling framework, sharing a common structure but differing in ways that provided leverage on our three main questions about the influence of temporal prolongnation and discontinuity.

Our general approach was to fit the free parameters of each model using the sequence of stimuli and the subject’s response on each trial and then to evaluate the optimized model performance using the behavioral assays described above. Where feasible, we derived analytic expressions for model performance on the behavioral assays. When analytic solutions were too complex or not possible to derive, we relied on Monte Carlo simulations. In this section, we provide general expressions for the model-fitting procedure and the predictions about the behavioral assays. In subsequent sections, we derive the specific equations used for each model and assay.

Model fitting was performed in the maximum likelihood framework. For each model, we found the parameter set *θ* that maximized the log-likelihood of observing the set of responses *R* given the set of stimulus sequences *S* across all trials. Let *x_i_* represent the evidence afforded by the *i*th stimulus presentation on trial *j*, and let *r_j_* represent the subject’s response (“low” or “high”) on that trial. In practice, we need to derive only the probability of one of the responses because *P*(*r_j_* = low) = 1 – *P*(*r_j_* = high). Assuming independence between trials, the log-likelihood of the data under each model is

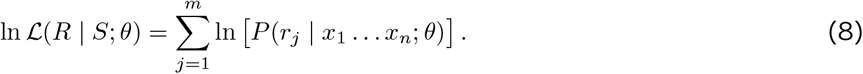

This function was numerically optimized for each model using the Nelder-Mead algorithm as implemented in scipy [58].

To obtain confidence intervals on the optimized log-likelihood and parameter values, we bootstrapped the maximum likelihood estimation procedure. On each of 1000 iterations, we randomly resampled trials (with replacement) before re-estimating the model parameters. The resampling procedure maintained the proportion of trials with each number of total samples. To obtain confidence intervals on the model parameter values, we used percentiles of the bootstrap distribution (i.e., for 95% CIs, we used the 2.5 and 97.5 percentile of the distribution).

We quantitatively compared the fit of the normative Linear Integration model to each of the other three candidates in terms of differences in the cross-validated log-likelihood (Δ*LL_Cv_*) of the choice data after optimizing the parameters. The Δ*LL_CV_* measure was defined so that positive scores indicated higher likelihoods for the Linear Integration model. To obtain cross-validated log-likelihood scores, we performed k-fold cross-validation by splitting the behavioral datasets by experimental session (k = 18). Aggregate Δ*LL_CV_* scores were calculated by summing the log-likelihoods from the model corresponding to each individual subject before taking the difference. Because the Linear Integration model is nested within the Leaky Integration model (they are equivalent if λ and *σ_∊_* both equal 0), we also used a likelihood ratio test to compare these models.

We further evaluated model performance in terms of how well model expressions for the three behavioral assays described in the previous section corresponded to the behavior of the subjects. Model expressions for the behavioral assays were derived using only the optimized model parameters and the generating statistics of the task, not the specific stimulus sequences on each trial.

In the following equations, *μ_x_* and *σ_x_* represent, respectively, the mean and standard deviation of the evidence distribution for the “high” condition in units of LLR. Note that the distributions for the two conditions are symmetric around LLR = 0, so the generating mean for the “low” condition is −*μ_x_*. Sometimes we will work only with the positive distribution corresponding to *μ_x_*, and at other times we will work with the set *D* = {−*μ_x_*, +*μ_x_*}. Finally, let *N* represent the distribution of sample counts across trials, and let *n* represent a specific count.

To obtain a model mPMF, we compute separate predictions for each generating distribution and sample count, and we then combine them into the full prediction using a weighted sum. Therefore, mPMF can be expressed, in general, as

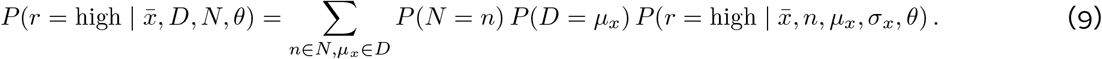

The model cPMF can be expressed, in general, as

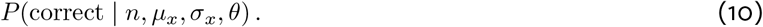

To generate a model RCF, we take the same approach as with the mPMF: we compute separate predictions for trials with different sample counts and then combine them into a full prediction. While the two psychometric functions represent probabilities, the reverse correlation represents a conditional expectation of stimulus evidence given response accuracy, *C*. Because we are conditioning on accuracy, and not choice, the RCF is defined in terms of evidence supporting the correct alternative, which we denote as 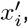 where 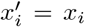 for the “high” distribution and 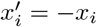 for the “low” distribution. Therefore, the RCF can be expressed, in general, as

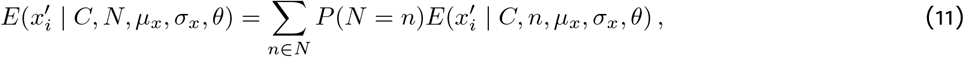

where *N* is the set of sample counts across trials, and *C* defines whether the RCF is for a correct or error response.

To drawatime-resolved RCF, we took a weighted average of these values across ordinal sample positions where the weights corresponded to the probability that a sample at time *t* was the *i*th sample in that trial. Probabilities were estimated empirically from the behavioral dataset.

In the following sections, we derive specific expressions for the forms of these functions under the assumptions of different models. Except where noted, the model predictions were computed using numerical integration algorithms as implemented in scipy.

### Linear Integration model

In the Linear Integration model, each stimulus affords a quantity of evidence towards the decision, which is noisily encoded, and the perceived evidence afforded by different samples are summed. The summed evidence represents a decision variable, *V*, which is defined in Equation 1. The response is determined by the sign of the decision variable at the end of the trial:

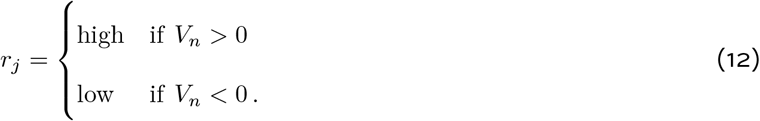

Assuming that the noise is independent for each sample, the decision variable *V* can be represented by a Gaussian distribution with a mean at the sum of the evidence, Σ *x_i_* and a standard deviation that grows with the square root of the number of samples, *n*. Integrating over the possible values of accumulated noise provides the probability of choosing each alternative (cf. equation 8):

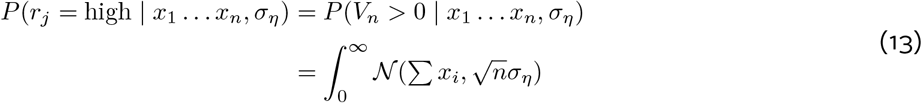

where 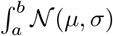 corresponds to the integral of a one-dimensional Gaussian distribution with mean *μ* and standard deviation *σ* taken with respect to the one-dimensional variable whose probability is defined by the distribution.

equation 13 shows that the choice probability given a particular sequence of samples depends only on the observation noise, number of samples, and mean evidence (*x̅* = ∑ *x_i_*/*n*). This establishes the mPMF as a key assay for comparison of behavior with the Linear Integration model:

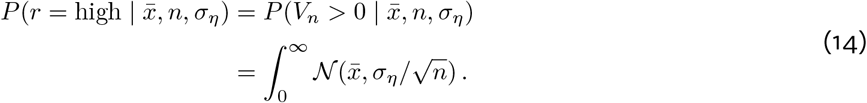

Because the cPMF is defined based on accuracy, it depends on both the stimulus distribution and the sensory noise. It can be computed from the expected distribution of decision variables. The mean of the decision variable distribution scales linearly with the number of samples, and its width is determined by summing the variance attributable to the random sampling of stimuli with the variance of the accumulated sensory noise (cf. equation 10):

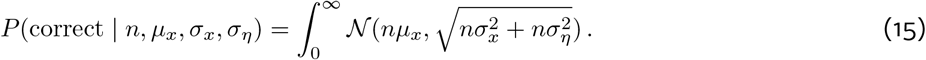

Under Linear Integration, the RCF will be flat across time for a given total number of samples. The height of the RCF conditional on a given sample count can be found by integrating over possible values of evidence and computing the conditional probability of accuracy on a trial featuring a sample with that value (cf. equation 11):

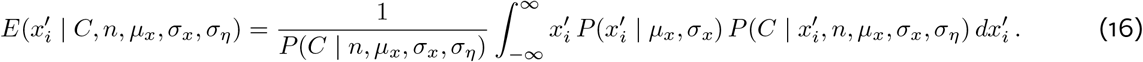

The first term in the right hand side of the equation is a normalization factor, which can be obtained from equation 15. The second term is a weighted integral of 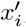, where the weights are defined by both the likelihood of observing sample 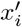 and the likelihood that a trial including that sample will lead to response *C*. The computation of this term depends on whether the trial has more than one sample. Single-sample predictions require marginalization only over sensory noise:

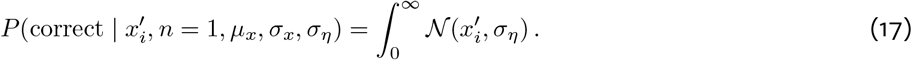

With multiple samples, however, it is necessary to marginalize over the contributions of sensory noise on sample *i*, *y_i_*, and the joint influence of evidence and noise on other samples:

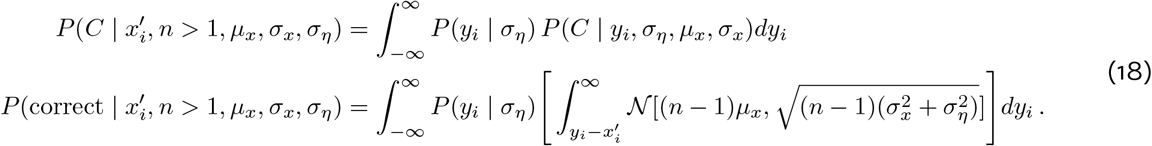

### Extrema Detection model

In the Extrema Detection model, the noisy perception of evidence afforded by each sample is compared against a threshold; if the threshold is exceeded, the process terminates in a commitment to the corresponding decision. Otherwise, the sample is discarded and the process continues. If the trial ends without a commitment, the process generates a random choice. The update equation that produces a decision variable is defined in Equation 2. The response is determined either by a commitment during the trial or by drawing a random response *g_j_* from *G* ~ Bernoulli(.5):

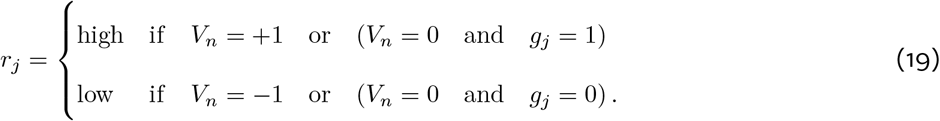

Unlike with Linear Integration, predicting the probability of choosing each alternative under the assumption of Extrema Detection requires knowledge about the specific sequence of stimuli. Intuitively, the prediction of the “high” choice is based on the probability that the noisy perception of a given sample, *X̂_i_* = *x_i_* + *ξ_η_*, will exceed the positive threshold weighted by the probability that the process has not yet terminated before sample *i* (cf. equation 8):

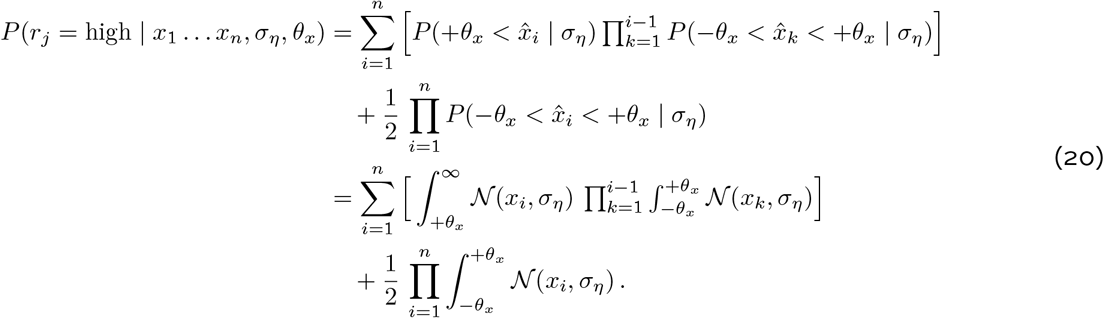

To predict the choice probability given a specific mean evidence value, *x̅*, it is necessary to marginalize over all possible stimulus sequences that could have produced that mean. Therefore, the basic approach is to compute (cf. equation 9):

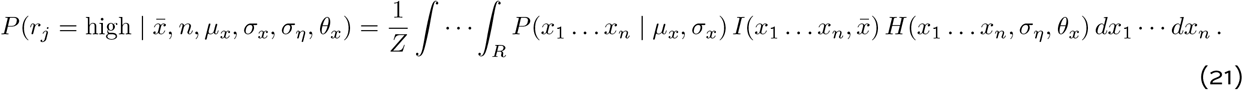

Working backwards, the function *H* gives the probability that the stimulus sequence *x*_1_ …*x_n_* will lead to a “high” response and can be computed as in equation 20. The function *I* is an indicator function that selects only those sequences with mean *x̅*. Non-zero response probabilities are further weighted by the probability of observing that sequence of stimuli, which is computed using the parameters of the evidence-generating distribution. Finally, the integrated values are divided by a normalizing constant:

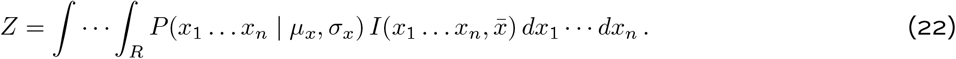

In practice, the discontinuity imposed by *I* requires us to approximate the integrals by sampling on a dense grid with bin width *ω*, so *I* becomes

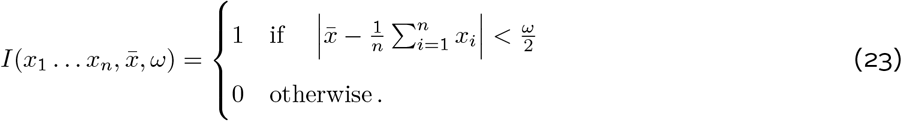

For predicting accuracy as a function of the number of samples, we take a similar approach as in equation 20 but use the generating statistics rather than the specific stimulus sequences (cf. equation 10):

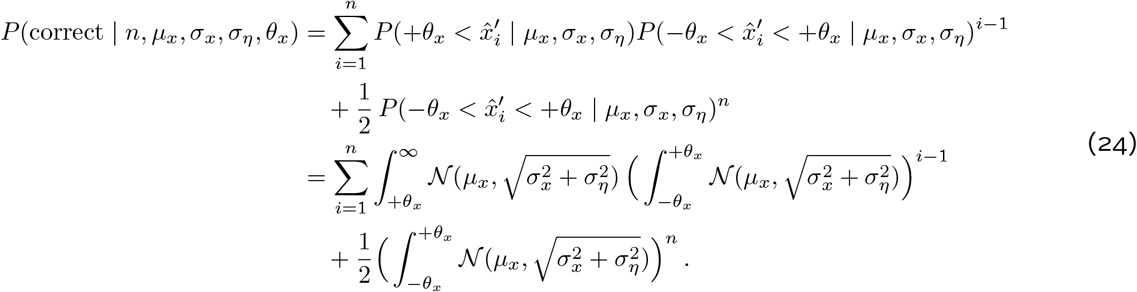

Predicting the reverse correlation function depends on the additional observation that the conditional distribution of evidence depends on whether or not the Extrema Detection process has terminated at that point. Let *T_i_* represent whether the process is still considering evidence at the ith sample, with *T_i_* = 0 representing an active sampling process and *T_i_* = 1 representing a terminated process. Once the process has terminated, the samples do not bear on choice; their conditional probability given the choice is determined by the generating distribution for the stimulus, with mean *μ_x_*. Further, because we are conditioning on accuracy, the active process must be sampling from a distribution that is truncated at −*θ_x_*. Therefore, the contributions of samples from these two distributions on sample *i* are weighted by the probability that the process has terminated prior to that sample (cf. equation 11):

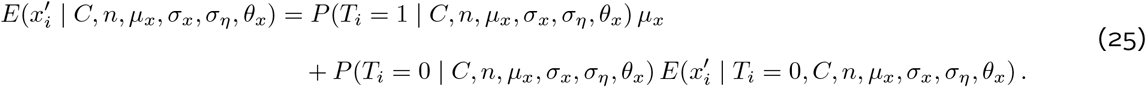

The conditional distribution for active processes can be further broken down by whether the noisily perceived evidence from a given sample, 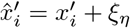, is above, below, or between the thresholds:

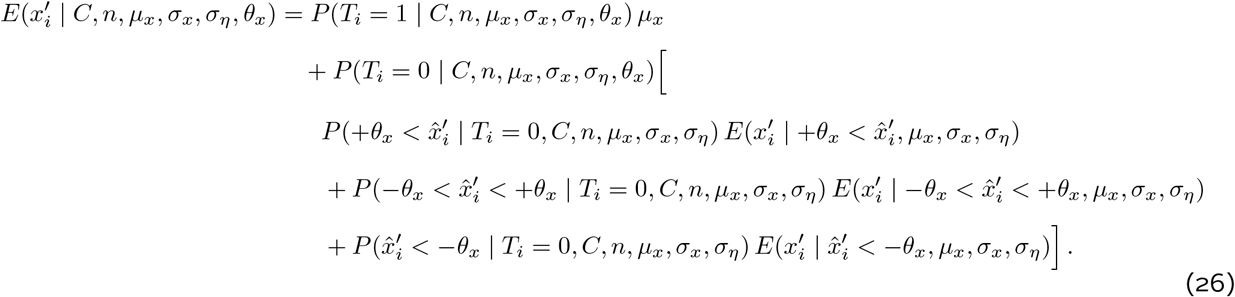

The conditional expectations in equation 26 require marginalizing over possible values of sensory noise:

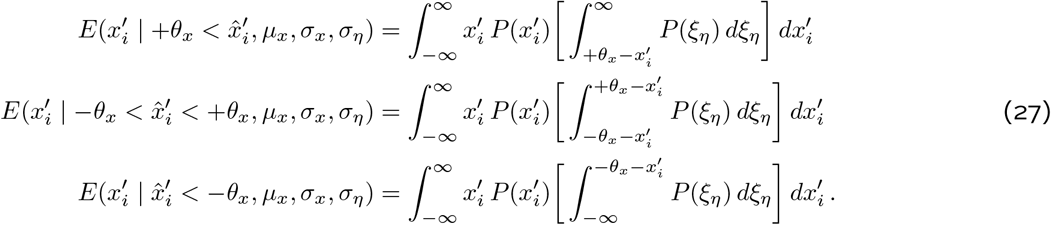

The conditional probabilities of observing evidence at different positions with respect to the threshold in equation 26 can be found using Bayes rule:

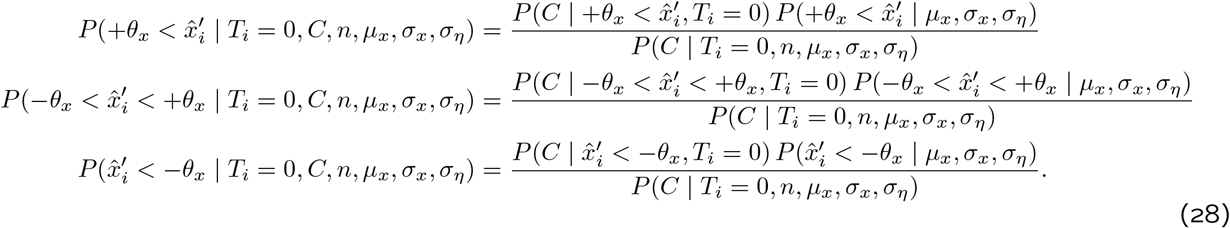

The conditional probability of accuracy given an observation between the thresholds in equation 28 can be found using equation 24 but substituting *n* – *i* for *n*. The conditional probability given observations above or below the threshold in equation 28 are given by the definition of the Extrema Detection process:

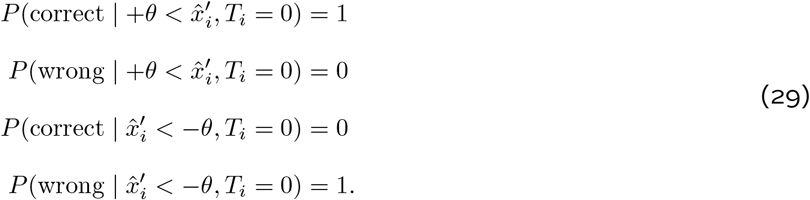

The second term in each numerator of equation 28 is the marginal probability of observing evidence in each segment, which depends on the parameters of the distribution that generates evidence and on the sensory noise:

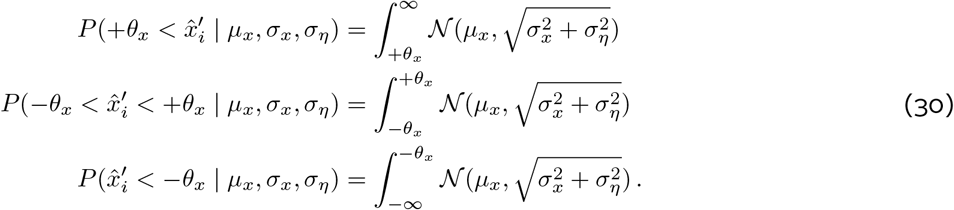

Each denominator in equation 28 is the probability of response accuracy conditional on the process not having terminated at sample *i*, which again can be found using equation 24 with *n* – *i* substituted for *n*.

So far, we have derived all the terms necessary for the calculation of the RCF (equation 26) except for the probability of having or not having terminated before sample i. A process that has not terminated prior to sample *i* has observed only intermediate values of noisy evidence up to that point, and the probability of this occurring can be computed using equation 28:

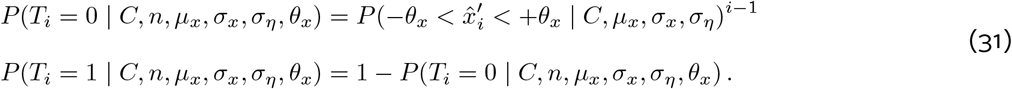

### Counting model

In the Counting model, each sample undergoes a transformation to a positive or negative increment, and a discrete count is maintained across samples. The process is defined in Equation 3. The response is determined by the sign of the decision variable at the end of the trial. Because *V* is maintained as a discrete count, it is possible that trials with an even number of samples will end in a tie (*V_n_* = 0). In this case, as with the Extrema Detection model, the response is generated using a random guess *g_j_* from *G* ~ Bernoulli(.5):

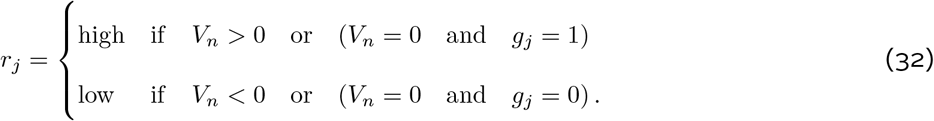

Therefore, the probability of a “high” response equals the sum of two probabilities: the probability of positive increments for more than half of samples or a positive increment in exactly half and a positive random guess. If we define 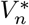 as representing the number of positive increments from *n* samples, the choice probability becomes (cf. equation 8)

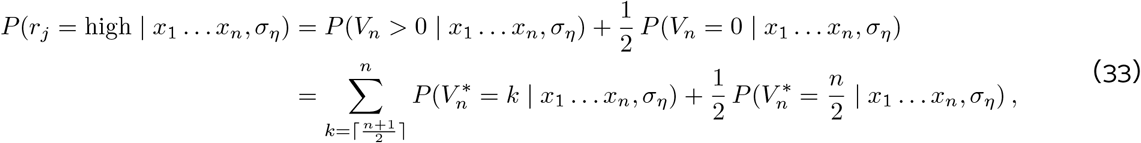

where ⌈.⌉ is the ceiling function. There is a unique probability of each sample producing a positive increment in the accumulator. For sample *x_i_* the probability of an increment is

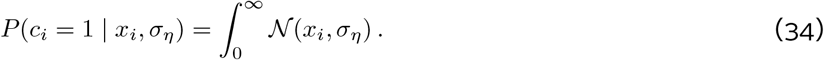

Further, observe that

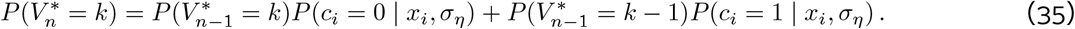

We can recursively apply this equation to compute the probabilities in equation 33.

Predicting choice probability from a specific mean evidence value (the mPMF) follows a similar approach to the one taken for the Extrema Detection model using equation 21. The only difference is the *H*(.) function; for the Counting model, we can use equation 33 to compute the choice probability for a given sequence of stimuli.

To generate a prediction for the cPMF under the Counting model, we also take a similar approach as in 33. But it is not necessary to consider each idiosyncratic sequence. Instead, the probability that the count of increments supporting the correct response takes any particular value is a Binomial random variable with success probability 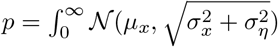. The sample psychometric function can therefore be expressed as (cf. equation 10)

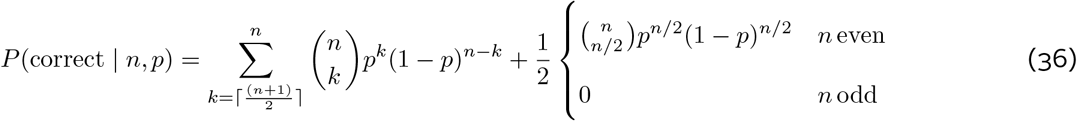

where 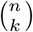 is the binomial coefficient function (*n* choose *k*). The first term on the right hand side of the equation gives the probability of success due to a count that favors the right choice, while the second term gives the probability of success due to guessing correctly after a tie.

The Counting model makes a qualitative prediction that accuracy on trials with an even number of samples will not exceed accuracy on trials with the next smallest odd number. This result follows from Equation 36, but the logic behind it can also be seen by considering the possible sequence of events on each trial. If the probability that a sample will support the correct choice is *p*, then the probability of responding correctly on a trial with two samples can be written as

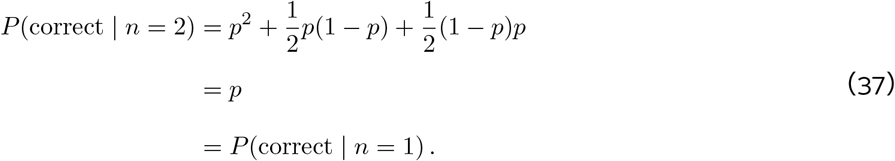

This shows the equivalence between accuracy on one-sample and two-sample trials. The logic can be extended to pairs of trial types with larger numbers of samples.

Similar to the case with Linear Integration, the RCF for the Counting model will be flat for a given total number of samples. It can be computed in general using equation 16 with a change in the conditional probabilities and normalization constant. The normalization constant can be found using equation 36. As with Linear Integration, the conditional probability of response accuracy given evidence on sample *i* is different for single and multiple sample trials. In fact, the two models make identical predictions for single sample trials (equation 17). For multi-sample trials, the conditional probability depends on whether sample *i* will produce a positive or negative increment in the count:

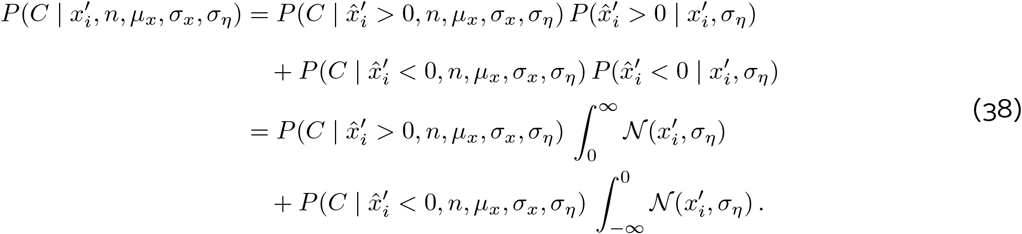

The conditional probabilities use the Binomial distribution following a similar logic to what we used when computing the cPMF (equation 36). If 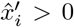, getting a correct response requires the remaining *n* – 1 samples to produce at least *n*/2 positive increments or *n*/2–1 positive increments and a lucky guess. If 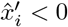, the remaining *n* – 1 samples must produce at least (*n* + 1)/2 positive increments or *n*/2 positive increments and a lucky guess.

### Leaky Integration model

The Leaky Integration model extends the Linear Integration model to account for two additional factors that could influence the decision variable as a function of time. The process is defined in Equation 4. As in Linear Integration, the response is determined by the sign of the decision variable at the end of the trial.

To fit the model, we use the fact that as Δ*t* → 0, the change in the decision variable during the gaps can be described with an Ornstein-Uhlenbeck (OU) process. If sample *i* at time *t_i_* is followed by gap duration 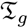, the OU process transforms the Gaussian distribution of the decision variable after sample 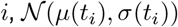, to another Gaussian distribution with the following mean and variance at the end of the gap:

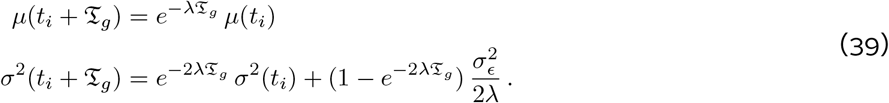

At the end of the trial, time *T*, the choice likelihood is given by integrating the positive density of the resulting distribution following all samples and gaps (cf. Eq. 8):

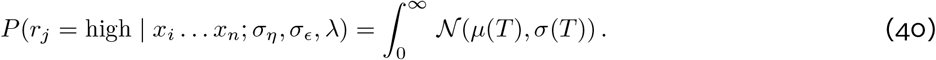

Analytic predictions for the behavioral assays could be obtained by extending the equations of the Linear Integration model for an OU process. However, the resulting equations would be complex because they would need to marginalize over all gap durations in the experiment. We avoid this complexity by using Monte Carlo simulation of model performance for the trials in the behavioral datasets (5 simulations per trial; Δ*t* = 100ms) and comparing the simulated model behavior to that of our subjects.

## Acknowledgements

RK is supported by the NIMH (R01-MH109180), the Simons Collaboration on the Global Brain (542997), and a Pew Scholarship in Biomedical Sciences. MLW is supported by the Simons Foundation as a Junior Fellow in the Simons Society of Fellows (527794).

---

**Figure S1:**
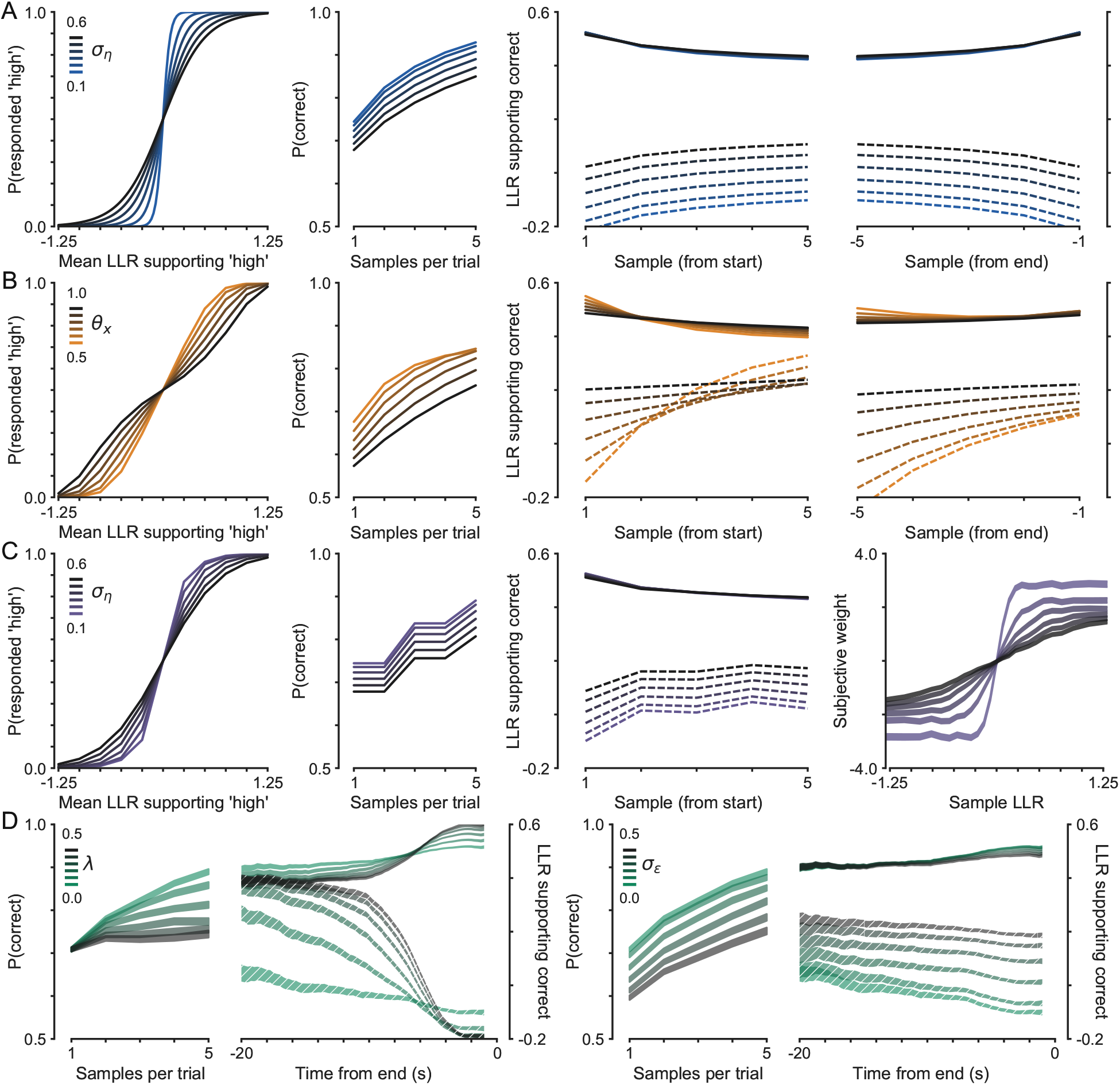
Computational model predictions for different behavioral assays. (A) Predictions of the Linear Integration model when varying the magnitude of sensory noise, *σ_η_*. (B) Predictions of the Extrema Detection model when varying the height of the detection threshold, *θ_x_*, while holding sensory noise constant (*σ_η_* = 0.2). (C) Predictions of the Counting model when varying the magnitude of sensory noise, *σ_η_*. (D) Simulated performance for the Leaky Integration model (N = 100,000 trials). The left panels show simulations varying the memory leak rate, *λ* while holding the noise terms constant (*σ_η_* = 0.4 and *σ_∊_* = 0). The right panels show simulations varying the memory noise magnitude, *σ_∊_* while holding sensory noise and memory leak constant (*σ_η_* = 0.4 and λ = 0). The widths of the bands show 95% confidence intervals.

**Figure S2:**
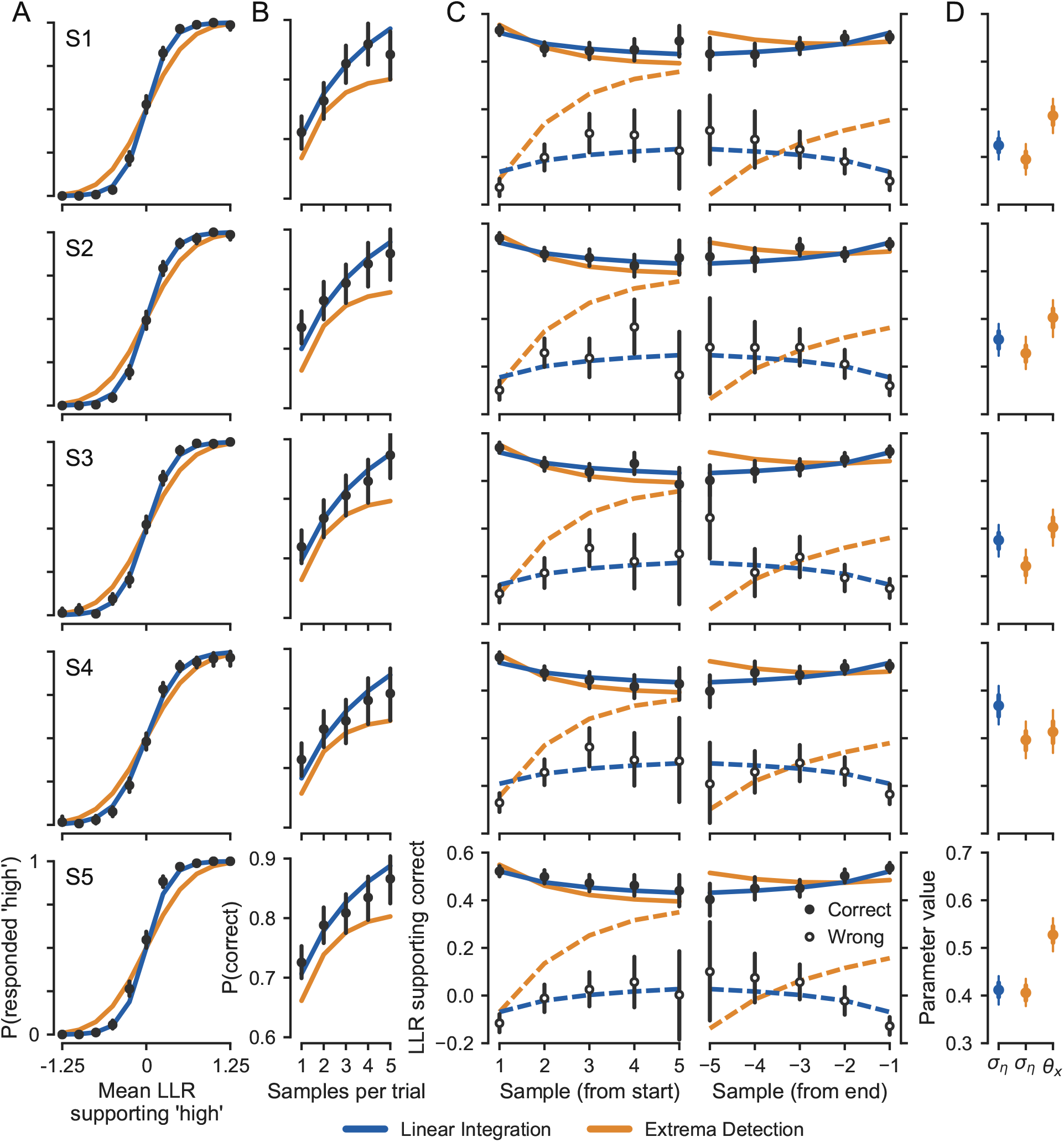
Decisions were based on integration of evidence across samples. Each row shows data and model fits for individual subjects. (A-C) Individual data and model fits for the three main behavioral assays, as in Figure 2A-C. (D) Estimated model parameters (points) and bootstrap confidence intervals (thick and thin error bars show 68% and 95% CIs, respectively).

**Figure S3:**
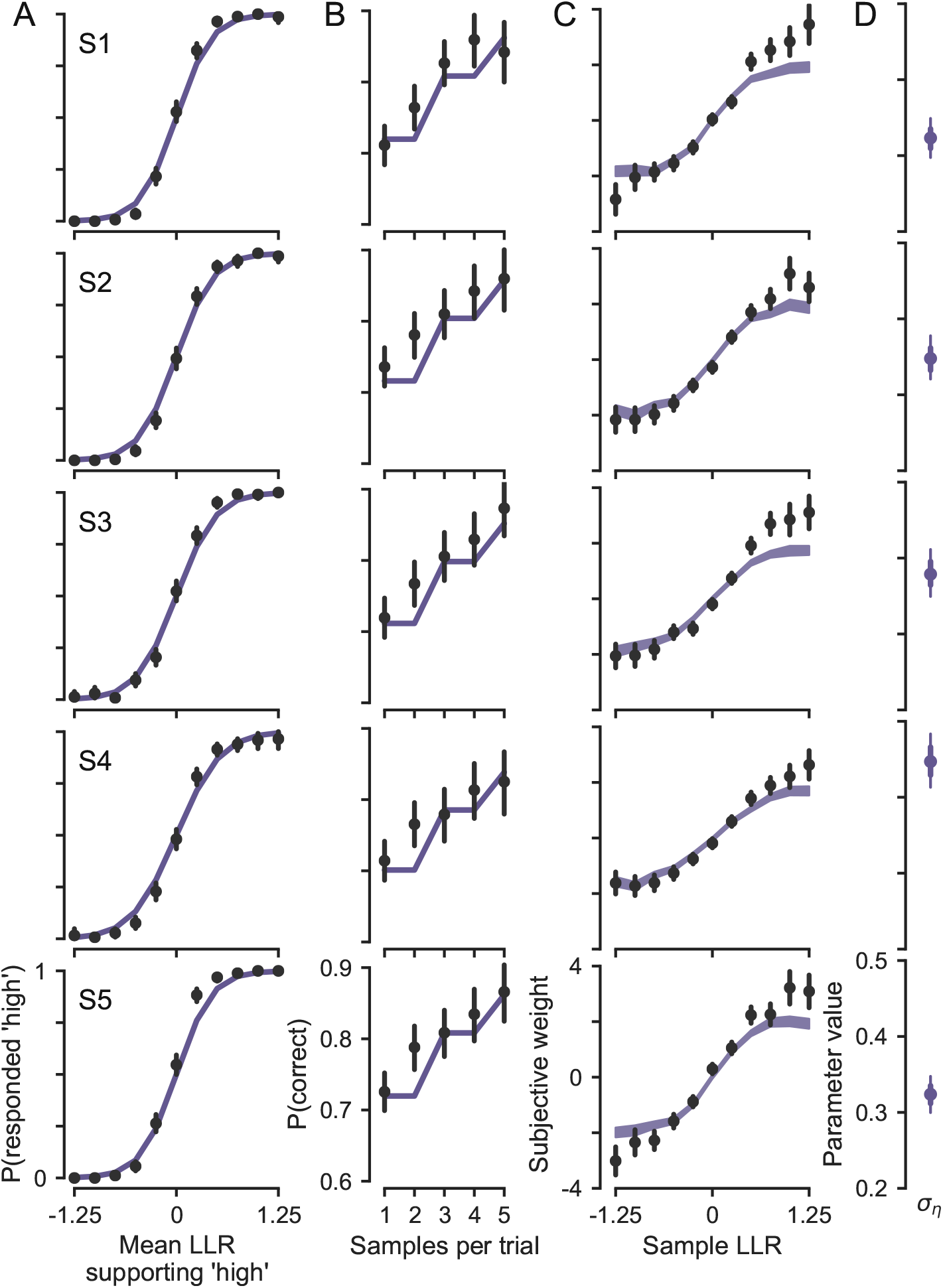
Integration of graded stimulus evidence. Each row shows data and model fits for individual subjects. (A) Individual data and model mPMFs. (B-C) Individual data and model fits corresponding to figure 3. (D) Estimated model parameters (points) and bootstrap confidence intervals (thick and thin error bars show 68% and 95% CIs, respectively).

**Figure S4:**
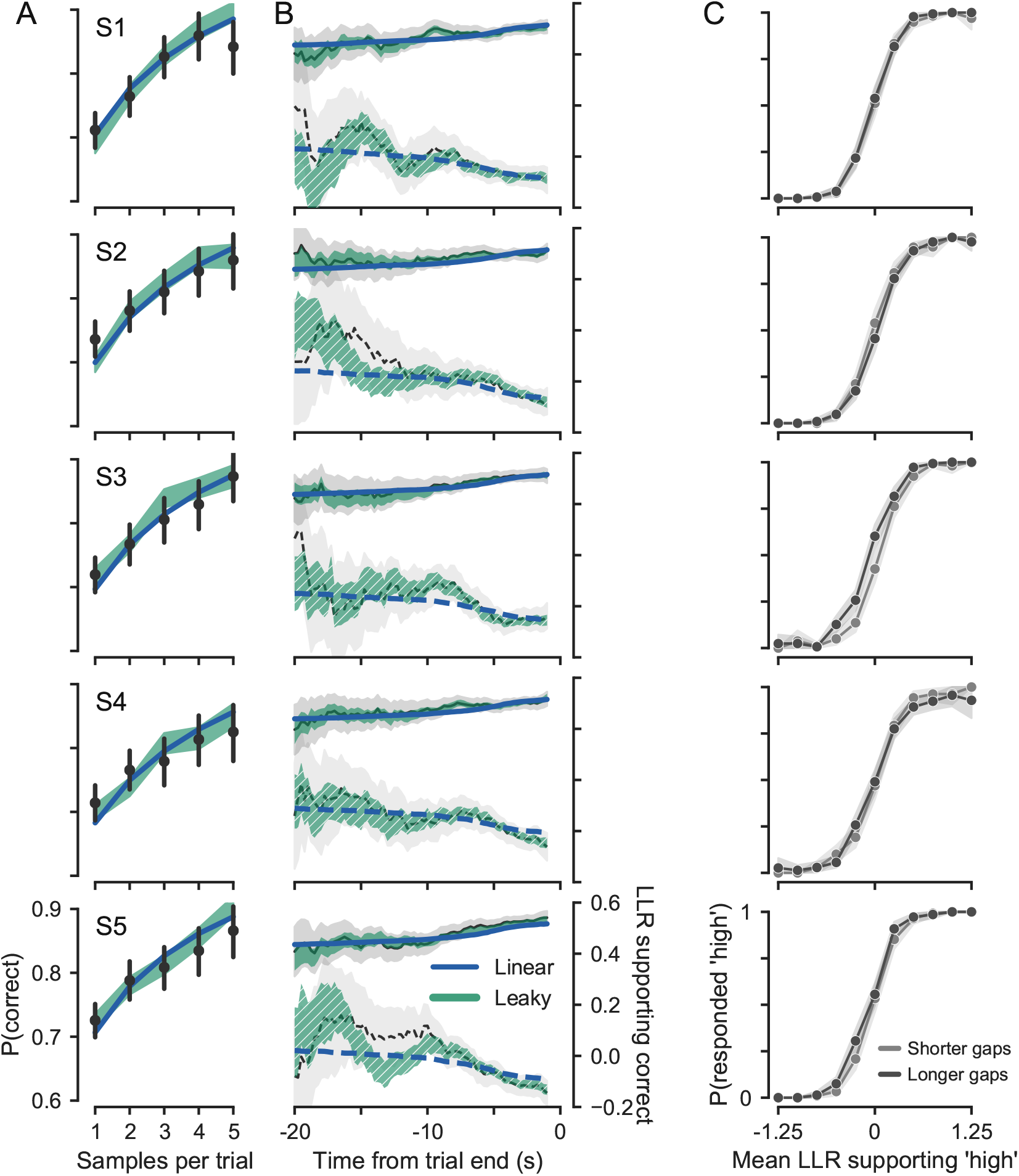
Minimal influence of memory leak or noise. Each row shows data and model fits for individual subjects. (A-C) Individual data and model fits corresponding to figure 4.

